# Melatonin drugs inhibit SARS-CoV-2 entry into the brain and virus-induced damage of cerebral small vessels

**DOI:** 10.1101/2021.12.30.474561

**Authors:** Erika Cecon, Daniela Fernandois, Nicolas Renault, Caio Fernando Ferreira Coelho, Jan Wenzel, Corentin Bedart, Charlotte Izabelle, Sarah Gallet Wimez, Sophie Le Poder, Bernard Klonjkowski, Markus Schwaninger, Vincent Prevot, Julie Dam, Ralf Jockers

## Abstract

COVID-19 is a complex disease with short- and long-term respiratory, inflammatory and neurological symptoms that are triggered by the infection with SARS-CoV-2. Invasion of the brain by SARS-CoV-2 has been observed in humans and is postulated to be involved in post COVID condition. Brain infection is particularly pronounced in the K18-*hACE2* mouse model of COVID-19. Here, we show that treatment of K18-*hACE2* mice with melatonin and two melatonin-derived marketed drugs, agomelatine and ramelteon, prevent SARS-CoV-2 entry in the brain thereby reducing virus-induced damage of small cerebral vessels, immune cell infiltration and brain inflammation. Brain entry of SARS-CoV-2 through endothelial cells is prevented by melatonin through allosteric binding to human angiotensin-converting enzyme 2 (ACE2), which interferes with the cell entry receptor function of ACE2 for SARS-CoV-2. Our findings open new perspectives for the repurposing of melatonergic drugs in the prevention of brain infection by SARS-CoV-2 and COVID-19-related long-term neurological symptoms.

## INTRODUCTION

As of 20 December 2021, the severe acute respiratory syndrome coronavirus 2 (SARS-CoV-2) is estimated to have infected globally approximately 270 Mio. people, including more than 5 Mio. deaths and more than 500,000 new confirmed infected cases per 24h, reported to WHO (https://covid19.who.int/). Although COVID-19, the disease caused by the SARS-CoV-2 virus, targets primarily the lungs leading to an acute respiratory disease, increasing evidence indicates a more widespread infection of other organs including the heart and blood vessels, kidneys, gut, and brain ^1^. Among the secondary symptoms, various neurologic signs and symptoms have been reported such as headache, nausea, anosmia, myalgia, hemorrhage, syncope, seizure, and stroke ^2^. Many of these symptoms remain long after the acute illness has passed, a phenomenon described as “long COVID”, post COVID-19 condition ^3^ or Post-Acute Sequelae of SARS-CoV-2 infection (PASC) ^4 5^. The estimated incidence of a neurological or psychiatric diagnosis in the following 6 months of infection is of 33% with no apparent correlation with severeness of COVID-19 in the acute phase, using electronic health records of over 236,000 patients with COVID-19 ^6^. Several possible mechanisms explaining the neurological symptoms and the neuro-invasive potential of SARS-CoV-2 have been discussed^7^. Mechanisms by which the virus accesses the brain include a direct viral infection of host endothelial cells, altering the tight junction proteins forming the blood-brain barrier (BBB) resulting in a leaky BBB (paracellular migration), or a phagocytosis by circulating immune cells followed by brain infiltration of these cells (“Trojan horse” strategy, by transcellular migration) ^7–9^. Another entry gate into the central nervous system is located at the neural-mucosa interface in the olfactory mucosa ^10^. Much attention has been given to identify potential treatments for lung infection and to control the cytokine storm, but only very few efforts to treat or prevent symptoms of post COVID condition have been made so far. Melatonin is a natural hormone produced by the pineal gland during the night with a wide range of effects on the central nervous system (CNS) including the regulation of the biological master clock in the hypothalamus and sleep on-set. Additional favorable effects are its neuroprotective, anti-inflammatory and anti-oxidant actions ^11^. Melatonin acts through a variety of target proteins ^12^ of which the two high-affinity G protein-coupled receptors, MT_1_ and MT_2_, are best-described ^13^. Currently marketed drugs (ramelteon, agomelatine, tasimelteon and slow-release melatonin) acting on MT_1_ and MT_2_ are indicated for insomnia, « jet-lag », and depression ^14^ and have been proven to show a good safety profile displaying few or no side-effects ^15, 16^. Recent systems-network-pharmacology studies revealed melatonin as one of the top-scoring molecules with potential anti-COVID-19 action ^17, 18^.

Based on this large spectrum of action of melatonin, its potential beneficial effects to treat COVID-19 have been postulated in several review articles ^19–22^. However, only very few experimental data from animal models or humans are currently available. We have recently shown that melatonin treatment of K18-*hACE2* mice (expressing the human ACE2 receptor) delayed the occurrence of severe clinical outcome and improved survival, associated with a dampening of virus-induced type I and type III interferon production in the lungs ^23^.

Here, we show that daily injection of melatonin and melatonergic compounds largely diminishes SARS-CoV-2 infection of the brain in the K18-*hACE2* COVID-19 mouse model, which displays high level of viral brain penetrance, by reducing viral entry through brain endothelial cells and damage of cerebral small vessels. This goes along with a concomitant reduction of neuroinflammatory markers and of markers of immune cell infiltration.

Furthermore, we identified a new binding target of melatonin, as melatonin reduces the entry of SARS-CoV-2 into brain cells by binding to ACE2, the SARS-CoV-2 cell entry receptor.

## RESULTS

### Melatonin treatment improves the clinical score and decreases viral load in the brain

To evaluate the potential beneficial effect of melatonin and the two clinically used melatonin receptor ligands agomelatine (AgoMLT) and ramelteon (RML) on SARS-CoV-2 infection in the brain, we chose the K18-*hACE2* mice, a robust model of brain infection by SARS-CoV-2^24, 25^. For melatonin, two doses were chosen, 10 mg/kg (MLT10) and 50 mg/kg (MLT50), aiming to maintain levels high over time as melatonin has a short plasma half-life of 20-30 minutes ^26^. Intra-peritoneal treatment of mice with compounds started two days before intranasal infection of mice and was repeated daily until sacrifice at day 7 post infection (DPI-7) at the latest. Lungs and brain were collected for biochemical and histological analysis. The course of development of COVID-19 in infected K18-*hACE2* mice was followed daily by evaluating body weight, activity and piloerection, respiration, lethargy and eye closure and a clinical score ranging from 0 to 14 was determined according to previously established guidelines with a score of 0 being perfectly healthy and a score of 14 being severely ill ^27^. Clinical scores worsened markedly from DPI-5 to DPI-6 in vehicle-treated animals, as expected, while a significant improvement on DPI-6 for MLT10 and MLT50 groups and a tendency for improvement for AgoMLT and RML was observed (**Fig. 1A**). At DPI-7 the beneficial effect was only maintained for MLT50 (**Fig. 1A**). This differential clinical evolution in treated groups is further evidenced when analyzing the frequency distribution of mice with high clinical score within each group, with a persistent beneficial effect at DPI-7, with the exception of MLT10 (**Fig. 1B**). These data suggest that treatment with melatonin and melatonergic drugs delay the onset of clinical symptoms and slow down disease progression. We then compared the viral load in the lungs and in the cerebral cortex by monitoring the transcripts for the SARS-CoV-2 nucleocapsid (or N-protein) by RT-PCR using a set of FDA-approved primers used to diagnose COVID-19 patients. In accordance with previous observations, the level of viral RNA in the lungs showed strong inter-individual heterogeneity in this model (**Fig. 1C**) ^23, 25^. Overall, the treatments did not seem to affect pulmonary viral load (**Fig. 1C**). The N-protein transcript was also detected in the cortex of 100 % of the mice in the vehicle group, although at overall levels approximately three orders of magnitude lower than those in the lungs (**Fig. 1D**). Despite the inter-individual heterogeneity, treatments with MLT10, AgoMLT and RML showed a tendency to lower viral load in the cortex, while MLT50 significantly decreases cortical viral load, an observation that was confirmed with two other FDA-approved N-protein primer pairs (**Fig. 1D**) (**Supplemental Figure S1A,B**). The effect of the treatment is confirmed when analyzing the distribution of mice into three categories according to their viral RNA load in the cortex classified per tercentiles (**Fig. 1E**). Whereas the majority of the mice in the vehicle-treated group falls into the category with the highest virus load, the treatment decreases the contingency in this category, and the frequency distribution was significantly different among the groups (Chi^2^-test, p=0.0065). When analyzing the contingency distribution of mice for each treatment compared to the vehicle group, MLT50 (Chi^2^-test, p=0.0357, **Supplemental Figure S1C**) was the most efficient treatment, as MLT50 largely suppressed the category of mice with the highest viral load and conversely increased the number of animals with low viral load in the cortex (**Figure 1E; Supplemental Figure S1C**). To a lower extent, MLT10 and RML also improved the distribution of mice (Chi^2^-test, p=0.0498 and 0.0357, respectively, **Supplemental Figure S1C**) increasing the intermediate category and decreasing the category of high cortical virus load. Decreased expression of the N-protein in treated mice was confirmed by microscope imaging of brain slices and the difference was even more pronounced in the heavily infected hypothalamic region of the brain (**Fig. 1F-G**) (**Supplemental Figure S2, 3**). These results indicate that treatment with melatonin drugs decreases infection of the brain by SARS-CoV-2, even in the K18-*hACE2* mouse model highly susceptible to brain infection.

**Figure 1:**
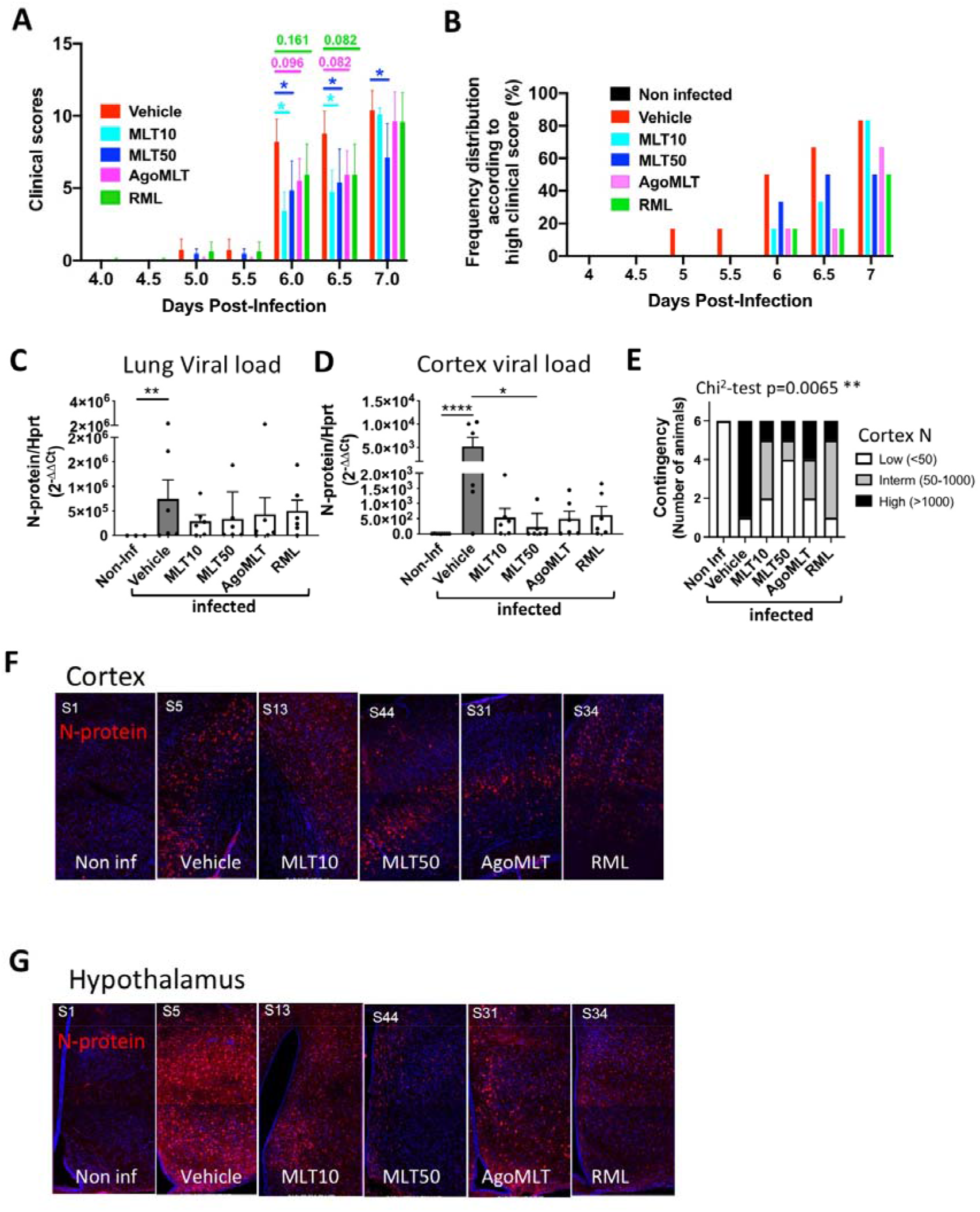
Treatment with melatonin receptor ligands improves clinical score and decreases viral load in the brain. **A.** Clinical score of infected mice. A clinical score was attributed to all the mice which were examined daily from post infection day 0 to day 4 or twice per day from day 4 to day 7. Mean clinical scores of mice for each group are shown. *p<0.05 by two-way ANOVA with two-stage linear step-up procedure of Benjamini, Krieger and Yekutieli as post-test for multiple comparisons. **B.** Frequency distribution (%) of infected mice with high clinical score (clinical score >6 until DPI-5.5 and >9 after DPI-6 for each group). **C-D.** RNA levels of viral N protein in the lungs (C) or in the cerebral cortex (D) of SARS-CoV-2 infected mice at DPI-7. *p<0.05, **p<0.01, ****p<0.0001 by Kruskal-Wallis test with two-stage linear step-up procedure of Benjamini, Krieger and Yekutieli as post-test for multiple comparisons. **E.** Contingency distribution of mice according to 3 groups of RNA level of viral N-protein in the cortex grouped by tercentiles (Chi2 test: p-value=0.0065). **F-G.** Immunolabeling for N-protein (red) and nucleus (blue) in the cortex (F) or hypothalamus (G) of K18-*hACE2* infected mice 7 days after SARS-CoV2 infection.

### Melatonin diminishes brain inflammation

The well-documented anti-inflammatory effect of melatonin has been hypothesized to be beneficial in COVID-19 treatment (see ^28^ for review). In cortex samples of SARS-CoV-2- infected K18-*hACE2* mice, MLT50 treatment tends to decrease the virus-induced mRNA expression levels of the proinflammatory cytokines *Tnf* (p=0.0514)*, Il1b* (p=0.1191) and *Il6* (p=0.0864) compared to vehicle-treated mice (**Fig. 2A-C**). Interestingly, the MLT50 treatment also tended to decrease mRNA levels of the chemokines *Cxcl1* (p=0.1607) and *Cxcl2* (p=0.0690) (**Fig. 2D-E**) and of markers of infiltrating macrophages *Trem1* (p=0.0964) and *Ms4a8a* (p=0.0223) (**Fig. 2F-G**). When considering the contingency distribution of infected mice in different treatment categories according to the tercentile classification of inflammatory markers in low, intermediate and high levels, a significant effect of MLT50 compared to vehicle was observed (**Fig. 2H-N**). MLT50 significantly decreased the number of mice with high levels and conversely increases the number of mice with low levels of *Il1b* (p=0.0384), *Il6* (p=0.0350), *Cxcl1* (p=0.0472), *Cxcl2* (p=0.0221), *Ms4a8a* (p-=0.0357) with tendencies for *Tnf* (p=0.0989) and *Trem1* (p=0.0578) (**Fig. 2H-N**). The other treatments (MLT10, AgoMLT and RML) had either no effect or a weaker effect than the MLT50 condition. Taken together, in particular MLT50 treatment had a favorable effect on preventing SARS-CoV-2-induced brain inflammation in K18-*hACE2* mice, in accordance with the lowest levels of viral brain infection observed in this group.

**Figure 2:**
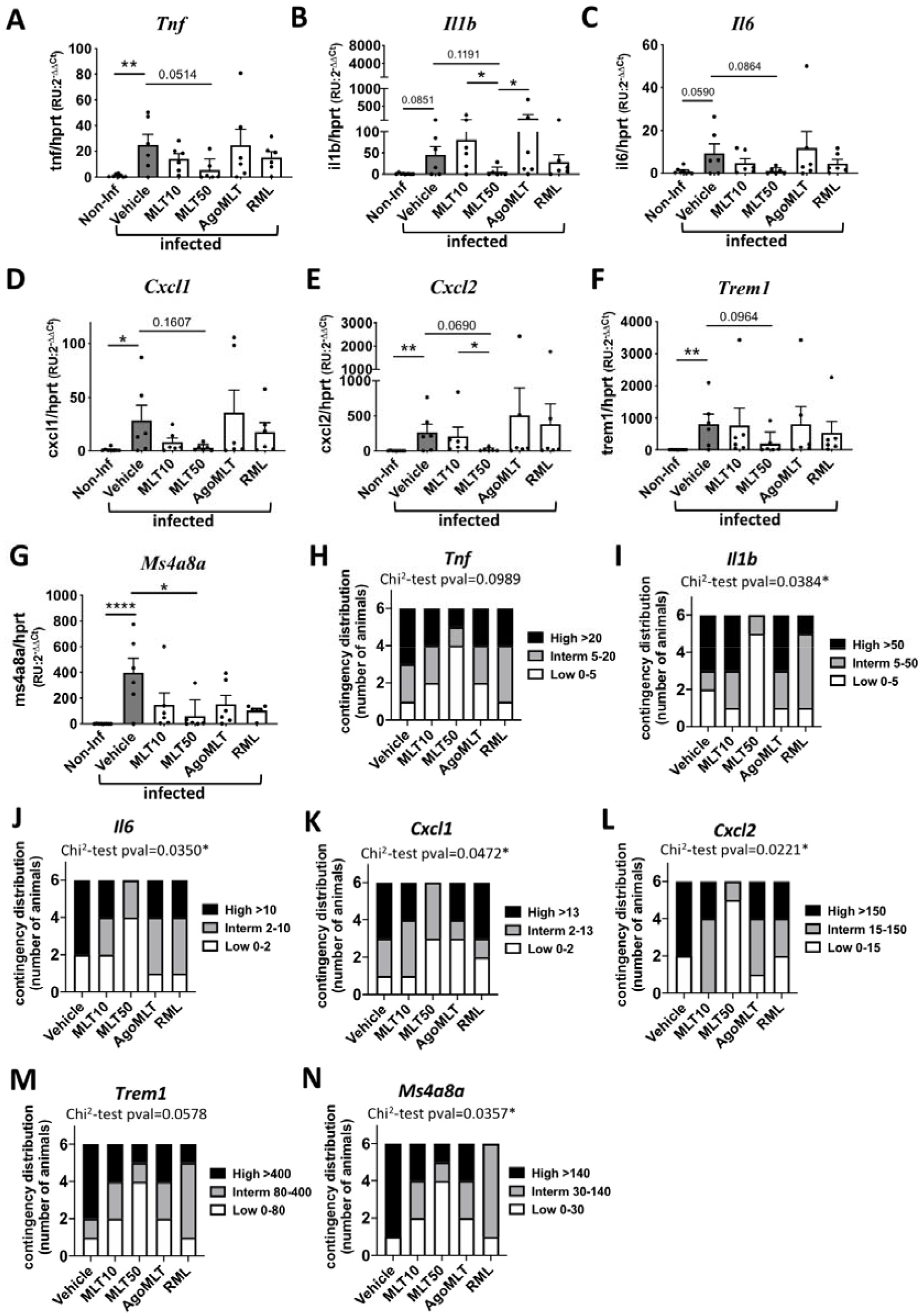
Melatonin diminishes brain inflammation. **A-G**. mRNA levels of *Tnf, Il1b, Il6, Cxcl1, Cxcl2, Trem1, Ms4a8a* in the cortex of SARS-CoV-2 infected mice at sacrifice day 7 were determined by RT-qPCR). *p<0.05, **p<0.01, ****p<0.0001, by Kruskal-Wallis test with two-stage linear step-up procedure of Benjamini, Krieger and Yekutieli as post-test for multiple comparisons. **H-N**. Contingency distribution of infected mice in different treatment categories depending on tercentile classification of inflammatory markers. p-values of Chi^2^-test between vehicle and MLT50 treatment considering “Low” and “High” levels of inflammatory markers are indicated (*p<0.05).

### Melatonin inhibits virus-induced damage of cerebral small vessels

Recent evidence indicated that COVID-19 damages cerebral small vessels in humans and K18-*hACE2* mice by infecting brain endothelial cells ^29^, which might favor brain infection by disrupting the blood-brain barrier. When staining cortical sections of our SARS-CoV-2-infected K18-*hACE2* mice for the basement membrane component collagen IV and the endothelial marker caveolin-1, we were able to replicate these results by showing the presence of string vessels that are formed when endothelial cells die (**Fig. 3A**). The analysis showed increased string vessel number and length upon SARS-CoV-2 infection (**Fig. 3B-C**).

**Figure 3:**
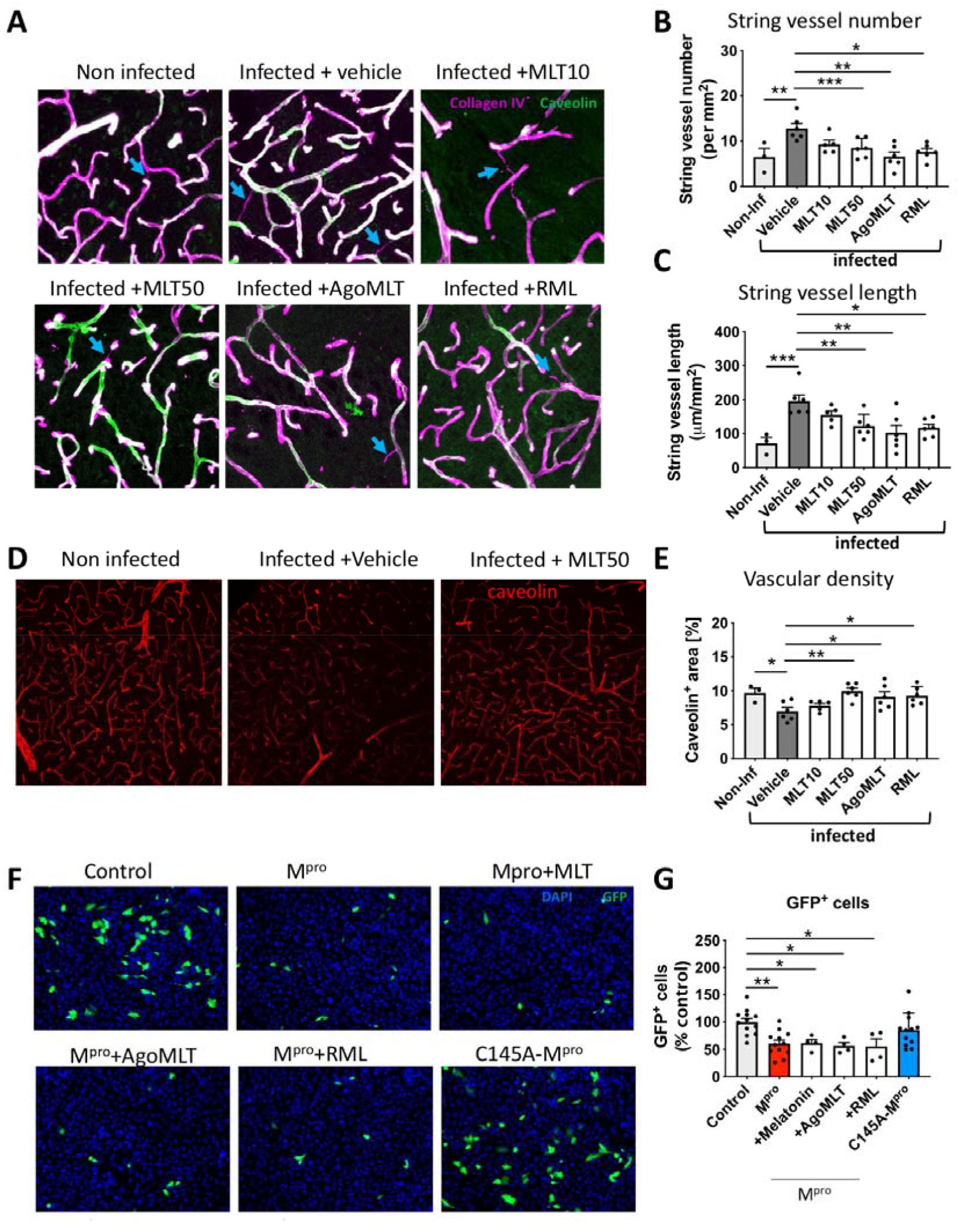
Melatonin inhibits SARS-CoV-2-induced damage of cerebral small vessels. **A**. Representative images of collagen IV (purple) and caveolin-1 (green) in the cortex of non-infected mice or infected mice and treated with either vehicle, MLT10, MLT50, AgoMLT or RML, 7 days post-infection with SARS-CoV-2. Empty basement membrane tubes, also known as string vessels are indicated with arrowheads. **B-C.** Quantification of string vessel number (**B**) and length (**C**) in non-infected mice and in treated SARS-CoV-2-infected mice**. D.** Caveolin staining of vascular system in brain slices of infected mice treated with vehicle or MLT50 versus non-infected mice. **E.** Quantification of caveolin positive area. **F.** Fluorescence images of human endothelial hCMEC/D3 cells transfected with GFP (control), GFP and M^pro^, or GFP and the inactive form C145A-M^pro^. A low number of hCMEC/D3 cells (GFP+) survived after expressing the M^pro^ protein while the expression of the inactive variant C145A-M^pro^ does not change the number of GFP+ cells. Treatment with MLT10, MLT50, AgoMLT, and RML did not interfere with the cell death-inducing effect of M^pro^. **G.** The numbers of surviving GFP+ hCMEC/D3 cells are plotted. *p<0.05, **p<0.01, ***p<0.001 by ordinary One-way Anova test followed by Dunnett’s post hoc test.

Interestingly, both parameters were significantly decreased in MLT50, AgoMLT and RML treated animals (**Fig. 3A-C**), while MLT10 showed only a tendency for decrease (**Fig. 3A-C**). Interestingly, the overall brain vascular density was found to be decreased in brains of mice infected with SARS-CoV-2, and MLT50 as well as RML and AgoMLT treatments protected the brains from capillary rarefaction (**Fig. 3D-E**). The mechanism by which SARS-CoV-2 damages cerebral small vessels involves the cleavage of the host protein NF-kappa-B

Essential Modulator (NEMO), shown to be required for the anti-viral immune response ^30^, by M^pro^, the main protease of SARS-CoV-2 ^29^. As melatonin has been shown to inhibit NF-kappa-B activation ^31, 32^, we evaluated whether melatonin and melatonergic ligands have an effect on M^pro^-induced cell death by investigating cell survival of cultured brain endothelial hCMEC/D3 cells transfected with a plasmid encoding for M^pro^. Treatment of hCMEC/D3 cells with melatonin, AgoMLT or RML (100µM, 48h) did not abrogate the effect of M^pro^ on cell death (**Fig. 3F-G**). Expression of the catalytically inactive C145A-M^pro^ mutant ^33^ was used as a control and was without effect on cell survival, as expected (**Fig. 3F-G**).

Collectively, these data indicate that all treatments prevent SARS-CoV-2-induced string vessel formation and protected the vascular system density. These effects could not be explained by inhibition of apoptosis of endothelial cells induced by the viral M^pro^ protease, suggesting thus that melatonin and its derivative act upstream of M^pro^, likely at the level of SARS-CoV-2 cell entry.

### Effect of melatonin on the expression of SARS-CoV-2 target proteins in the brain

To explore an action of melatonin in endothelial cells occurring upstream of the M^pro^ viral protease activity, such as SARS-CoV-2 cell entry mechanisms, we analyzed the effect of the treatments on the expression levels of the major components of the SARS-CoV-2 entry machinery: ACE2, neuropilin 1 (NRP1) and TMPRSS2 ^34, 35^. SARS-CoV-2 infection or compound treatments did not change the global mRNA levels of human *ACE2* (under the control of the K18 promoter)*, Nrp1* or *Tmprss2* neither in the cortex or in the lungs (**Supplemental Figure S4**). Similarly, no effect was observed on the expression of mouse *Ace2*, which does not participate in the virus entry in this model but could indicate an effect of the ligands on the regulation of the endogenous ACE2 promoter (**Supplemental Figure S4**). Knowing that the expression of melatonin receptors can be restricted to specific cell subpopulations in the mouse brain ^36^, we analyzed their expression on brain slices of the cortical vasculature of mice by RNAscope. Expression of MT_1_ and MT_2_ melatonin receptors (encoded by the *Mtnr1a* and *Mtnr1b* genes, respectively) were specifically expressed in collagen IV-positive endothelial cells in the brain (**Fig. 4A**). These results indicate that vascular endothelia might be the target of melatonin and its derivatives mediating their protective effects against brain infection.

**Figure 4:**
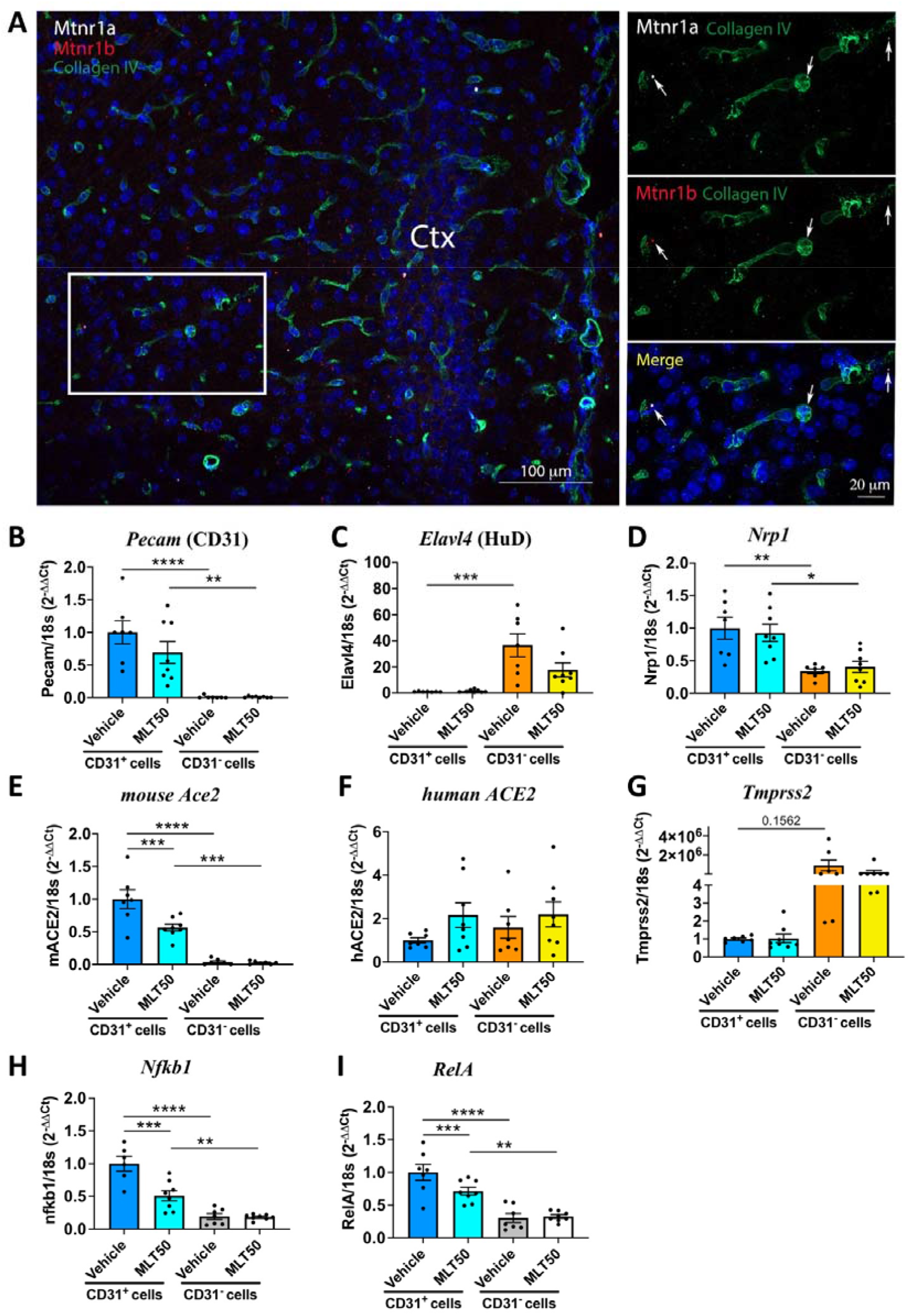
Effect of melatonin on the expression of components involved in cell entry of SARS-CoV-2 in the brain. **A**. Detection in mouse brain of melatonin receptor transcripts, *Mtnr1a* (white) and *Mtnr1b* (red), in the cortical vasculature (positive for collagen IV). **B-I**. mRNA levels of *Pecam* (CD31, endothelial marker), *Elavl4* (HuD, neuronal marker), *Nrp1*, mouse *Ace2*, human *ACE2*, *Tmprss2*, *Nfkb1* and *RelA* in sorted cortical cells positive or negative for CD31 from non-infected K18-*hACE2* mice. *p<0.05, **p<0.01, ***p<0.001, ****p<0.0001 by One-way Anova with post-hoc Tukey test for correction of multiple comparison.

We next isolated the CD31^+^ endothelial cell population by FACS sorting from the cortex of *K18-hACE2* mice treated with vehicle or MLT50 for 9 days to further evaluate the effect of melatonin on gene expression. Successful enrichment of endothelial cells in the CD31^+^ fraction *vs.* neurons in the CD31^-^ fraction was confirmed by the strong reciprocal expression of *Pecam* (CD31) in CD31^+^ cells and *Elavl4* (HuD, neuronal marker) in CD31^-^ cells (**Fig. 4B-C**). Consistent with the notion that brain endothelial cells are SARS-CoV-2 entry cells, expression of *Nrp1* and murin *Ace2*, two SARS-CoV-2 entry receptors, was particularly prominent in CD31^+^ cells with an enrichment of 3 and 34 times, respectively, over CD31^-^ cells (**Fig. 4D-E**). Expression of human *ACE2* (under the control of the K18 promoter) was similar in both fractions (**Fig. 4F**), while *Tmprss2* was predominantly expressed in CD31^-^ cells (**Fig. 4G**).

*Nrp1* levels were unaffected by MLT50 treatment while murine *Ace2* expression was decreased by half in CD31^+^ cells from MLT50-treated mice (**Fig. 4D-E**). MLT50 treatment did not modify the expression of the human *ACE2* under the control of the K18 promoter (**Fig. 4F**). Additionally, the MLT50 treatment downregulated *Nfkb1* and *RelA* expression, two genes of the NFκB pathway (**Fig. 4H-I**), the decrease of which might participate in the reduction of endothelial inflammation.

Collectively, these results show that brain endothelial cells are potential melatonin target cells for receptor-mediated melatonin effects and that melatonin decreases the expression of the endogenous ACE2 and NFκB pathway genes in these cells thus potentially contributing to the protective effect of melatonin on cerebral small vessels. However, downregulation of mouse *Ace2* cannot explain the reduction of virus entry in *K18-hACE2* mice, which is dependent on human ACE2 expressed under the control of the K18 promoter and the latter is not downregulated by melatonin, suggesting another inhibitory mechanism by melatonin.

### Binding of melatonin to ACE2 interferes with SARS-CoV-2 entry into brain endothelial cells

To address whether melatonin interferes directly with viral entry into host cells we used a SARS-CoV-2 pseudovirus luciferase reporter assay in HEK293 cells. Naïve HEK293 cells are resistant to infection but become highly susceptible for infection upon ectopic expression of human ACE2 (**Fig. 5A**). Melatonin inhibited ACE2-dependent virus entry in a concentration-dependent manner, with an estimated EC_50_ of 20 µM and a maximal inhibition of 60-70% at the highest concentration (**Fig. 5B**). These results were recapitulated in human brain endothelial hCMEC/D3 cells, in which pseudovirus cell entry is also shown to be dependent on the level of ACE2, and melatonin significantly inhibited the entry of the SARS-CoV-2 pseudovirus by 40% in hCMEC/D3 cells (**Fig. 5C**). These data show that melatonin directly interferes with ACE2-dependent SARS-CoV-2 entry.

**Figure 5:**
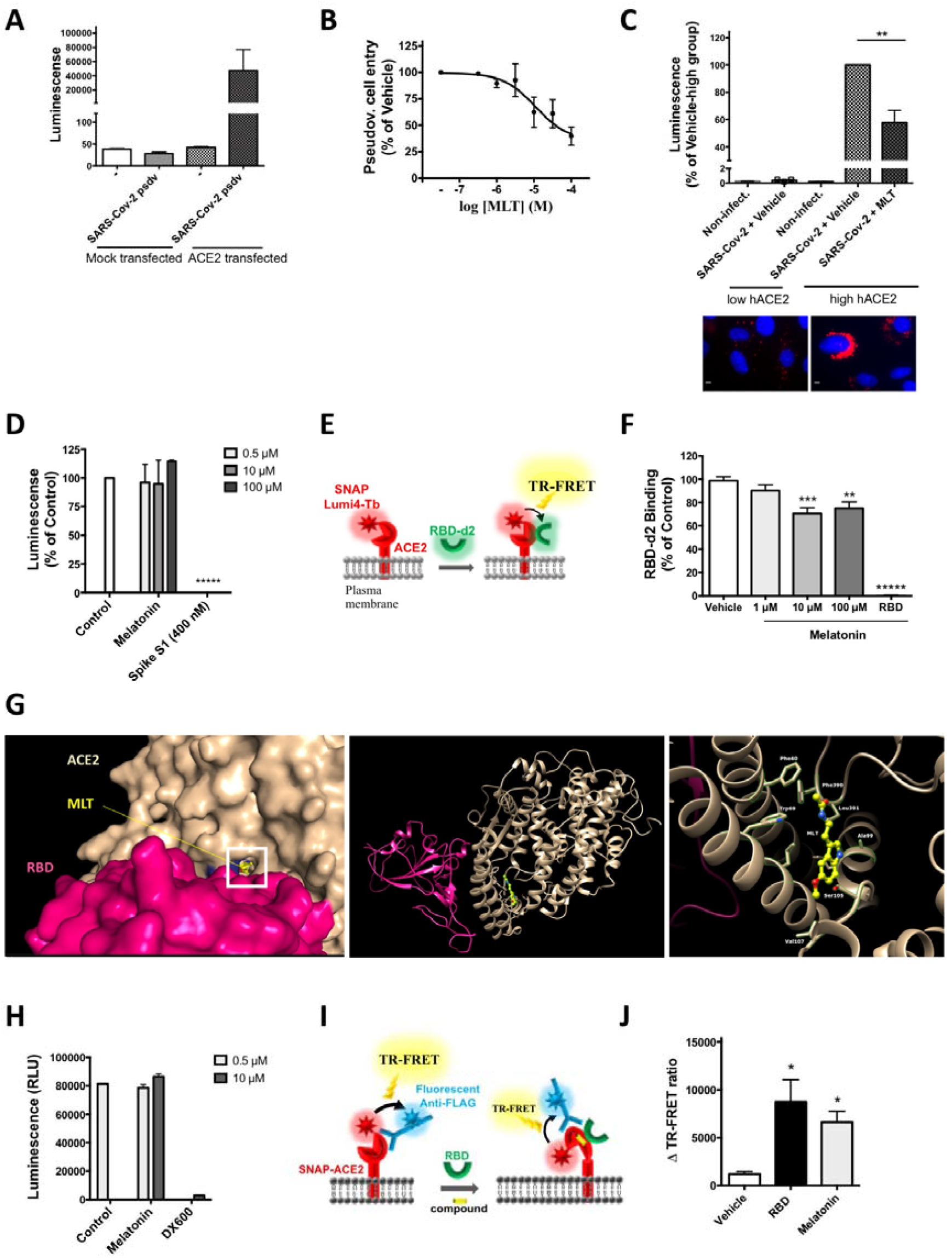
Melatonin inhibits SARS-CoV-2-cell entry by interfering with the spike-ACE2 interaction. **A.** SARS-CoV-2 cell entry assessed by pseudovirus luciferase reporter assay in HEK293 cells transfected with a human ACE2 expression plasmid or mock transfected. Data are expressed as mean ± SEM of 3 independent experiments, each performed in duplicates. **B.** Concentration-response curves of melatonin on SARS-CoV-2 pseudovirus cell entry in HEK293 cells expressing human ACE2. Data are expressed as mean ± SEM of 3 independent experiments, each performed in triplicates. **C.** Inhibition of SARS-CoV-2 pseudovirus cell entry by melatonin (100µM) in hCMEC/D3 cells transduced with human ACE2. Data are expressed as mean ± SEM of 3 independent experiments, each performed in triplicates. Bottom panel show ACE2 expression in transduced and non-transduced cells by immunofluorescence (red = ACE2; blue = DAPI-stained nucleus; scale bar = 10 µm). **D.** In vitro binding assay of biotinylated S1 spike protein (50 nM) to immobilized human ACE2 in the absence or presence of melatonin at 0.5 10 and 100 μ; a competition by an excess of non-labelled S1 (400 nM) determines the specific signal. **E**. Scheme illustrating the TR-FRET-based binding assay of RBD-d2 to SNAP-tagged ACE2 labelled with Lumi4-Tb at the cell surface. **F.** Change in TR-FRET signal of RBD-d2 (5 nM) binding to Lumi4-Tb-SNAP-ACE2 by increasing concentrations of melatonin. Competition by an excess of non-labelled RBD (300 nM) determines the specific signal. Data are expressed as mean ± SEM of 3 independent experiments, each performed in triplicates. **G**. Docking and molecular dynamic-based prediction of melatonin binding to the crystal structure of ACE2 in complex with the RBD domain of SARS-CoV-2 virus. At t0 (left panel), the molecular docking of melatonin into ACE2/RBD complex (6M17) shows a unique entry channel for melatonin located in close proximity to the interaction surface between ACE2 and the viral spike protein (left panel). At t=500 ns of simulation using glycosylated ACE2/RBD complex (6VW1), a global view of the complex shows the migration of MLT to the interior side of two ACE2 helices interfacing RBD (central and right panel). Most hydrophobic interactions are exhibited in a zoomed view centered onto the binding site (right panel). **H.** ACE2 enzyme activity in the absence or presence of melatonin at 0.5 and 10 μ; DW600 inhibitor (10 μ) is shown as a positive control. **I.** Scheme illustrating the assay probing conformational change within ACE2 based on intramolecular TR-FRET between Lumi4-Tb-labelled SNAP-ACE2 and d2-labelled anti-FLAG tag antibody. **J.** ACE2 conformational change in HEK293 cells in the presence of non-labelled RBD (5 nM), or melatonin (100 µM). Data are expressed as change of TR-FRET signal (Δ TRFRET ± SEM) of 4 independent experiments, each performed in triplicates. *p<0.05; **p<0.01: ***p<0.005: *****p<0.0005 by one-way ANOVA, followed by Dunnett’s multiple comparisons test compared to vehicle group.

We then determined the effect of melatonin on the spike/ACE2 interaction in an in vitro assay measuring the binding of recombinant biotinylated receptor binding domain (RBD) of spike to purified ACE2 immobilized on the plate. Melatonin was unable to interfere with RBD binding to ACE2 whereas an excess (400 nM) of non-biotinylated S1 significantly diminished RBD binding, as expected, indicating that melatonin does not directly compete with RBD binding to ACE2 (**Fig. 5D**). We then employed a second, time-resolved FRET (TR-FRET)-based cellular spike-ACE2 binding assay ^37^. The assay measures the binding of RBD to ACE2 in a cellular context, by following the energy transfer between the N-terminal SNAP-tagged human ACE2 labelled with the energy donor terbium (Tb) and RBD labelled with the energy acceptor d2 (RBD-d2) (**Fig. 5E**). The energy transfer reports on the molecular proximity between the two proteins coupled to the energy donor and acceptor if they are within a maximal distance of less than 10 nm ^38^. Binding of the RBD-d2 tracer was readily detectable in this assay and was partially inhibited by micromolar concentrations of melatonin with maximal inhibition levels reaching 20 to 30%, as compared to an excess of unlabeled RBD defining the specific TR-FRET signal (**Fig. 5F**). This reduction in TR-FRET signal can be either due to a direct competition of RBD binding to ACE2 by melatonin or to an allosteric effect of melatonin modifying the conformation of the RBD-ACE2 complex that moves the energy donor and acceptor apart. To address the question of whether melatonin modifies the affinity of RBD for ACE2 we performed competition TR-FRET binding assays with increasing concentration of unlabeled RBD in the absence and presence of melatonin. The pIC_50_ value of 7.80 ± 0.06 (n=3) confirmed the high-affinity binding of RBD which was, however, not modified by 10 or 100 µM of melatonin (pIC_50_ 7.77 ± 0.05, and 7.59 ± 0.08, respectively; n=3, data not shown). Taken together, the partial inhibition of viral entry and RBD binding to ACE2 by melatonin and the absence of any competitive response of melatonin on the RBD-ACE2 interaction strongly suggests an allosteric behavior and conformational modifications within the RBD-ACE2 complex induced by melatonin.

To gain insights in the potential binding mode of melatonin to the RBD-ACE2 complex, we performed a computational blind docking of melatonin using an exploration grid comprising the entire RBD and the same volume counterpart of ACE2 of three solved crystal structures of RBD-ACE2 complex, 6M17, 6M0J and 6VW1 ^39, 40^. None of the docking solutions revealed the existence of a melatonin binding pocket on RBD. The top-ranked solution revealed a previously unknown melatonin binding cavity on ACE2 that was located in close proximity to but not directly at the RBD-ACE2 interface (**Fig. 5G**). Molecular dynamics simulations confirmed the melatonin binding site depicting the stability and the specificity of melatonin binding to ACE2 in the N-glycosylated 6VW1 PDB structure through three replicates of 500 ns (**Supplemental Figure S5**). The stable binding site identified in the replicates, R1 and R2, is located close to the two helices of ACE2 (helix 19-52 and helix 53-89) interacting with RBD. Melatonin binds into a site delimited by Phe40, Trp69, Leu73, Ala99, N-glycoslyated Asn103 and Lys562 as observed in two of the three replicates. The third replicate shows a rapid dissociation of melatonin from 100 ns. This new binding site is distinct from the central catalytic site of ACE2, an observation which is consistent with the absence of any effect of melatonin on the enzyme activity of ACE2, as assessed in an *in vitro* enzymatic assay (**Fig. 5H**).

The location of the binding site close to the two helices of ACE2 interacting with RBD indicates that melatonin might modulate the conformation of the ACE2 interface thus impacting on the spike-ACE2 interaction. To test this hypothesis experimentally, we developed an intra-molecular TR-FRET assay to probe the movement of ACE helix 19-52 by measuring the distance between a Flag-tag introduced in front of ACE helix 19-52, and the SNAP-tag of ACE2 (**Fig. 5I**). As expected, unlabeled RBD, which modifies the position of ACE2 helix 19-52 compared to the apo-ACE2 state, changed the TR-FRET signal, reflecting RBD-induced conformational changes in ACE2 (**Fig. 5J**). Melatonin also triggered a change in TR-FRET signal, confirming its binding to ACE2 and its impact on the position of ACE2 helix 19-52 (**Fig. 5J**).

Taken together, our data indicate that melatonin binds to an allosteric binding site at ACE2 and modulates the RBD-ACE2 complex by modifying the position of helix ACE2 19-52 helix that contacts RBD, directly impacting on viral cell entry.

## DISCUSSION

SARS-CoV-2 infects several tissues including the brain, an observation which has been largely neglected as considered irrelevant for the acute phase of COVID-19. The description of persisting symptoms, several months after the acute phase, like increased incidence of headache, fatigue and neurological symptoms, point to a potential role of the brain as target of the virus. Strategies to protect the brain might be therefore of interest to prevent long-term consequences of SARS-CoV-2 infection. We show here in the K18-*hACE2* COVID-19 mouse model that treatment with melatonin, specially at a high dose, protects the brain by inhibiting SARS-CoV-2 entry into this organ. Melatonin binds to an allosteric binding site at ACE2, discovered in this study, that is connected to the molecular interface interacting with the viral spike protein thus preventing ACE2 to serve as an efficient SARS-CoV-2 entry receptor.

Additional beneficial effects of melatonin may arise from the fact that melatonin drives endothelial cells to a less reactive inflammatory state due to lower expression of NFkB pathway genes. Additionally, reduced endogenous expression of *Ace2* could also participate in the decrease of viral entry in models where endogenous receptors are sensitive to SARS-CoV-2. We identified brain endothelial cells as the possible main melatonin target cells expressing ACE2 and MT_1_ and MT_2_ receptors. Our data indicate that melatonin has a major effect on the protection of cerebral small vessels by preventing SARS-CoV-2 entry, SARS-CoV-2-induced string vessel formation and reduction in vascular density. The protective effect of melatonin and its clinically used derivatives in K18-*hACE2* mice, a model of high SARS-CoV-2 infection compared to humans, puts them in a privileged position as repurposing candidates for long COVID studies in humans.

Several reviews suggest the beneficial use of melatonin to treat COVID-19 contrasts with the only few experimental evidences ^19, 22, 41^, a scenario that prompted us to evaluate the overall effect of melatonin and agomelatine and ramelteon, two highly potent clinically-approved melatonin analogs. Despite the fact that all three compounds act on high-affinity melatonin receptors ^13, 42^, their effects were not identical in our study. Clearest effects were observed in the MLT50 group, followed by AgoMLT and RML, and finally MLT10. The difference between MLT10 and MLT50 argues rather against the involvement of high-affinity receptors which should be saturated at both doses. The similar pattern of effects observed for MLT50, AgoMLT, and RML for most parameters would suggest the involvement of a common, most likely low-affinity target, which could be ACE2. The need for the higher dose of melatonin (MLT50) compared to AgoMLT and RML could be explained by the shorter plasma half-life of melatonin compared to the synthetic drugs. The superior effect of MLT50 compared to AgoMLT and RML in many parameters could be either a dose effect or due to additional anti-oxidant properties of melatonin at this high dose that are not observed with AgoMLT and RML ^11^.

We identified brain endothelial cells as the possible main melatonin target cells expressing ACE2 and MT_1_ and MT_2_ receptors. A growing body of evidence indicate the susceptibility of endothelial cells to be infected by SARS-CoV-2 ^43, 44^ in a ACE2-dependent manner, contributing to microvascular pathology observed in the brains of SARS-CoV-2-infected patients ^29, 44^. MT_1_ receptor expression was observed in human cerebral mircovessels ^45^ and activation of MT_1_ and MT_2_ receptors in smooth muscle cells to mediate vasoconstriction and vasodilation, respectively ^42^. In rat microvasculature, melatonin acts on endothelial melatonin receptors to reduce acute inflammation by inhibiting leukocyte rolling and adhesion ^46, 47^. A potential role of endothelial melatonin receptors in the modulation of vascular tone has been also suggested ^48^. Our complementary data indicate that melatonin, most likely by targeting high-affinity melatonin receptors in brain endothelial cells, might have additional beneficial effects in the context of COVID-19 by decreasing the expression of ACE2 and NFκB pathway genes.

The tissue-specific efficacy of melatonin treatment was striking, from no significant effect on viral load in the lungs to a substantial blockage of infection in the brain. Several factors might underlie this tissue selectivity of the melatonin effect. Firstly, the SARS-CoV-2 infection follows a well-characterized dynamic, with the airway and lungs being the primary affected tissues while systemic infection is more tardive. Although we cannot discard potential beneficial effects of melatonin treatment at earlier stages, the absence of any effect of melatonin on the overall lung pathology argues against this possibility at least in the K18-*hACE2* model ^23^. Secondly, and most importantly, the mechanisms of virus invasion significantly differ between the first phase and the systemic spreading. In the early phase of infection, the upper and lower airway epithelia and the lung alveoli are the primary cells targeted by SARS-CoV-2 and are also the location where active viral replication takes place ^49, 50^. High virion concentration together with high expression of ACE2 in these cells ^51^, is certainly a favorable condition for virus cell entry, which is difficult to counteract by melatonin. In contrast, the systemic spreading of the virus infection is characterized by relatively low viremia and low expression of ACE2 in the case of the brain tissue in humans ^52^, conditions more favorable to an effect of melatonin.

Two main mechanisms through which SARS-CoV-2 might reach brain tissues have been proposed and involves either a neuronal route from nerve terminals in contact with high viral load at the nasal cavity, or though systemic viremia and BBB crossing. The later might occur through direct viral invasion of host endothelial cells lining the vascular wall (transcellular migration), through brain infiltration of infected immune-system cells (“Trojan horse” strategy) or through disrupted BBB or caused by the virus infection (paracellular migration) ^7–9^. Our data provide evidence for the two first mechanism. The prevention of SARS-CoV-2- induced damage of cerebral small vessels argues in favor of the transcellular migration route.

The low levels of markers of infiltrated macrophages in the brain observed in the MLT50 group suggests that melatonin might also prevent the “Trojan horse” strategy of virus brain invasion. Of note, high melatonin concentration has been previously shown to decrease the expression of cell-adhesion molecules necessary for cell migration through the endothelial cell layer also in a receptor-independent mechanism ^32, 53^. The disclosed mechanisms of action of melatonin in interfering with the spike/ACE2 interaction and with cell migration at high concentrations probably explains the need for in vivo administration of high melatonin dose and the tissue-specificity of the protective effect. This conclusion is in line with the inhibitory effect of indole chloropyridinyl esters on SARS-CoV-2 infection ^54^ and of melatonin and other indoles on Swine coronavirus cellular entry ^55^. However, the underlying mechanism of these in vitro effects observed at super high, millimolar, melatonin concentrations, remained unclear.

The therapeutic potential of melatonin in COVID-19 is supported by the observation that melatonin users show a 52% reduced likelihood of positive SARS-CoV-2 PCR test in African Americans ^56^. Several clinical trials based on melatonin treatment were conducted during the sanitary crisis and the results of some of them have been gradually released and have underscored a beneficial effect of melatonin use on sleep quality and outcomes of COVID-19 patients ^57^. Among them, although with a small number of patients, a randomized, double-blind clinical trial, concluded on the efficacy of melatonin as an adjunctive therapy in hospitalized patients with COVID-19, improving clinical symptoms and contributing to a faster recovery ^58^. In addition, a randomized clinical trial designed to assess the efficacy of melatonin resulted in lower thrombosis, sepsis, and mortality rate in COVID-19 patients ^59^. Finally, treatment with fluvoxamine, one of the mechanisms of which is to increase plasma melatonin levels, has reduced the need for hospitalization in high-risk ambulatory patients with COVID-19 ^60^. Altogether, these first clinical results highlight a therapeutic potential of melatonin in COVID-19. Considering the reduction in brain infection and the decrease in the central inflammatory response with melatonin treatment, it is predicted that long-term neurological disorders could be avoided/attenuated with melatonin treatment. Melatonin is a medicine that can be obtained without a prescription. Unfortunately, retrospective studies, seeking to determine the efficacy of drugs against neurological disorders in COVID-19, and which would be based on cohorts of already existing patients taking or not taking melatonin supplements, do not allow a well-controlled analysis because the dose and the time of taking melatonin-derived medicines are usually not known. In addition, the selection of melatonin patient group included in the retrospective analyzes may be initially biased as patients with previous neurological disorders (fatigue, depression, memory loss…) are more inclined to use alternative supplements such as melatonin. Thus, an analysis directly devoted to studying the effects of melatonin treatment, specially at high doses, in relieving long-term neurological effects would be required.

When considering to use melatonin, agomelatine or ramelteon in the context of COVID-19, the chronobiological and sleep initiating effects of these three molecules have to be managed^14^. Application at the right time, in the evening before bedtime, and for a restricted period (probably during and shortly after the acute phase of infection), will minimize the risk of dysregulation of the circadian system and of sleep induction. Additional strategies include the chemical optimization of these melatonergic compounds to improve the specificity and affinity for ACE2 vs. MT_1_ and MT_2_ receptors.

Our study describes the first allosteric binding site of ACE2 and melatonin is the first allosteric modulator of ACE2 which negatively regulates ACE2 binding to the SARS-CoV-2 spike protein. This opens additional opportunities for the development of ACE2-specific drugs that interfere with the spike interaction ^61^. Importantly, binding of melatonin to this allosteric binding site of ACE2 did not interfere with ACE2 enzyme activity thus avoiding potential severe side effects as ACE2 activity is essential for the proper function of the renin-angiotensin system.

In conclusion, we disclosed ACE2 as a new binding target of melatonin, which leads to allosteric negative modulation of the spike/ACE2 interaction. This effect on the SARS-CoV-2 entry receptor function of ACE2, combined with the inhibition of NFKB expression and endogenous ACE2 expression in endothelial cells, are most likely the predominant mechanisms for impaired SARS-CoV-2 brain invasion by melatonin and its derivates.

## METHODS

### Animals

K18-*hACE2* C57BL/6 transgenic mice (males, 10-week-old), which expresses human ACE2 driven by a human cytokeratin 18 (K18) promoter (Jackson Laboratory, https://www.jax.org/strain/034860) were housed in an animal facility of biosafety level 3 (BSL3) at the French National Veterinary School in Maisons–Alfort, with water and food ad libitum. All animal experiments were approved by the ANSES/EnvA/UPEC Ethics Committee (CE2A-16) and authorized by the French ministry of Research under the number APAFIS#25384-2020041515287655 v6, in accordance with the French and European regulations.

### SARS-CoV-2 virus infection

At day of infection (day post-infection DPI-0), mice were infected via intra-nasal inoculation of SARS-CoV-2 (10 µL each nostril, 10^4^ TCID_50_ in total) in Dulbecco’s modified Eagle medium, under isoflurane anesthesia. The SARS-CoV-2 strain *BetaCoV/France/IDF/200107/20* was supplied by the Urgent Response to Biological Threats (CIBU) hosted by Institut Pasteur (Paris, France) and headed by Dr. Jean-Claude Manuguerra. The human sample, from which the strain *BetaCoV/France/IDF/200107/2020* was isolated, has been provided by Dr O. Paccoud from the La Pitié-Salpétrière Hospital (Paris, France).

### *In vivo* treatment with melatonergic compounds

Melatonin, agomelatine and ramelteon were purchased from abcr GmbH (Karlsruhe, Germany). Compounds were reconstituted in vehicle solution (5% ethanol in sterile saline solution). K18-*hACE2* transgenic mice were randomly divided into the following groups (6 mice/group): vehicle, melatonin 10mg/kg (MLT10), melatonin 50mg/kg (MLT50), Ramelteon (RML, 10 mg/kg), and Agomelatine (AgoMLT, 20 mg/kg), and were intraperitoneally (i.p.) injected with vehicle or melatonergic ligands daily, one hour before lights off (to minimize the risk of disturbing the natural daily rhythm of endogenous melatonin production). The treatment started 2 days before virus inoculation and continued until the end of the experiment (7 days post-infection, DPI-7). A group of non-infected mice was housed under the same conditions and similarly monitored during the whole experiment. Mice were uniquely identified using ear tags and provided an acclimation period of 1 week before the experiment. Mice were supplied nutrient gel when weights began to decrease. Mice that met the human endpoint criteria were euthanized to limit suffering. The study was ended at DPI-7 and surviving mice were sacrificed at that time for comparative analysis. Lung and brain samples were taken directly after sacrifice and stored at -80°C until analysis.

### Clinical score evaluation

All mice were weighed and examined for the presence of clinical symptoms daily, or twice per day after DPI-4, when symptoms were evident. The score for clinical symptoms was attributed following an IACUC approved clinical scoring system, and included the following criteria: body weight, posture/fur, activity/ mobility, eye closure, respiratory rate. Scoring was performed according to standard guidelines ^27^ with a maximal score of 14. Mice died either naturally from the disease or were sacrificed for ethical reasons when reaching a clinical score of 5 for 2 parameters and for 2 consecutive observation periods, or if weight loss was equal to or greater than 20%.

### Isolation of endothelial cells by fluorescence-activated cell sorting (FACS)

15 non-infected K18-*hACE2* male mice were injected ip either with vehicle (ethanol 5% in saline) or melatonin 50mg/Kg (MLT50) daily for 9 consecutive days. The morning of the 10^th^ day, brain was extracted and cortex immediately submerged in the cellular dissociation buffer.

Mice cortex was enzymatically dissociated using a Papain Dissociation System (Worthington, Lakewood, NJ) to obtain a single-cell suspension. Cells were incubated with blocking Purified Rat Anti-Mouse CD16/CD32 (553142, Mouse BD Fc Block™) for 15 min at 37°C before incubation with primary antibodies Alexa Fluor 647 Rat Anti-mouse CD31 (BD Pharmigen cat:563608) for CD31 cell labeling and Alexa Fluor 647 Rat IgG2a,k Isotype control (BD Pharmigen cat:557690). Antibody labeling was performed for 1h at 37°C. All steps of labeling and washing were performed in PBS-BSA 2%. Labeled cells were suspended in 5% glucose Hanks’ Balanced Salt Solution before sorting. FACS was performed using an ARIA SORP cell sorter cytometer device (BD Bioscience, Inc). The sorting parameter were based on measurements of Alexa Fluor 647 fluorescence (excitation: 633nm; detection: bandpass 670/30nm) by comparing cell suspensions from non-labeled cortex of K18-*hACE2* mice and the corresponding Isotype control. For each animal 4,000 CD31-positive and 4,000 tomato-negative cells were sorted directly into 10 μl of lysis buffer (0.1% Triton® X-100 and 0.4 unit/μl RNaseOUT™ (Life Technologies)).

### RNA extraction and real time qPCR analyses

For infected mice, total RNA was extracted from cortex by using the ReliaPrep™ FFPE Total RNA Miniprep System (cat: Z1002, Promega), homogenization was carried out in a glass-glass tissue grinder. RNA samples were immediately quantified in a NanoDrop One (Thermo Scientific v2) and stored at -80°C until the retro-transcription step. For gene expression analyses, total RNA was reverse transcribed using High capacity cDNA Reverse transcription kit (Applied Biosystems™ ref 4368814). For sorted CD31 positive cells, a linear preamplification step was performed using the TaqMan® PreAmp Master Mix Kit protocol (Applied Biosystems™ ref 4488593). Real-time PCR was carried out using TaqMan™ Universal Master Mix II (Applied Biosystems™ ref 4440049) on the Applied Biosystems 7900HT Fast Real-Time PCR System. For gene expression the TaqMan® probes listed below were used (Table 1). SARS-CoV-2 viral infection was assessed by the CDC 2019-Novel Coronavirus Real-Time RT-PCR Diagnostic Panel as described elsewhere ^62, 63^. All gene expression data were analyzed by the 2^-ΔΔCt^ method.

**Table 1.**
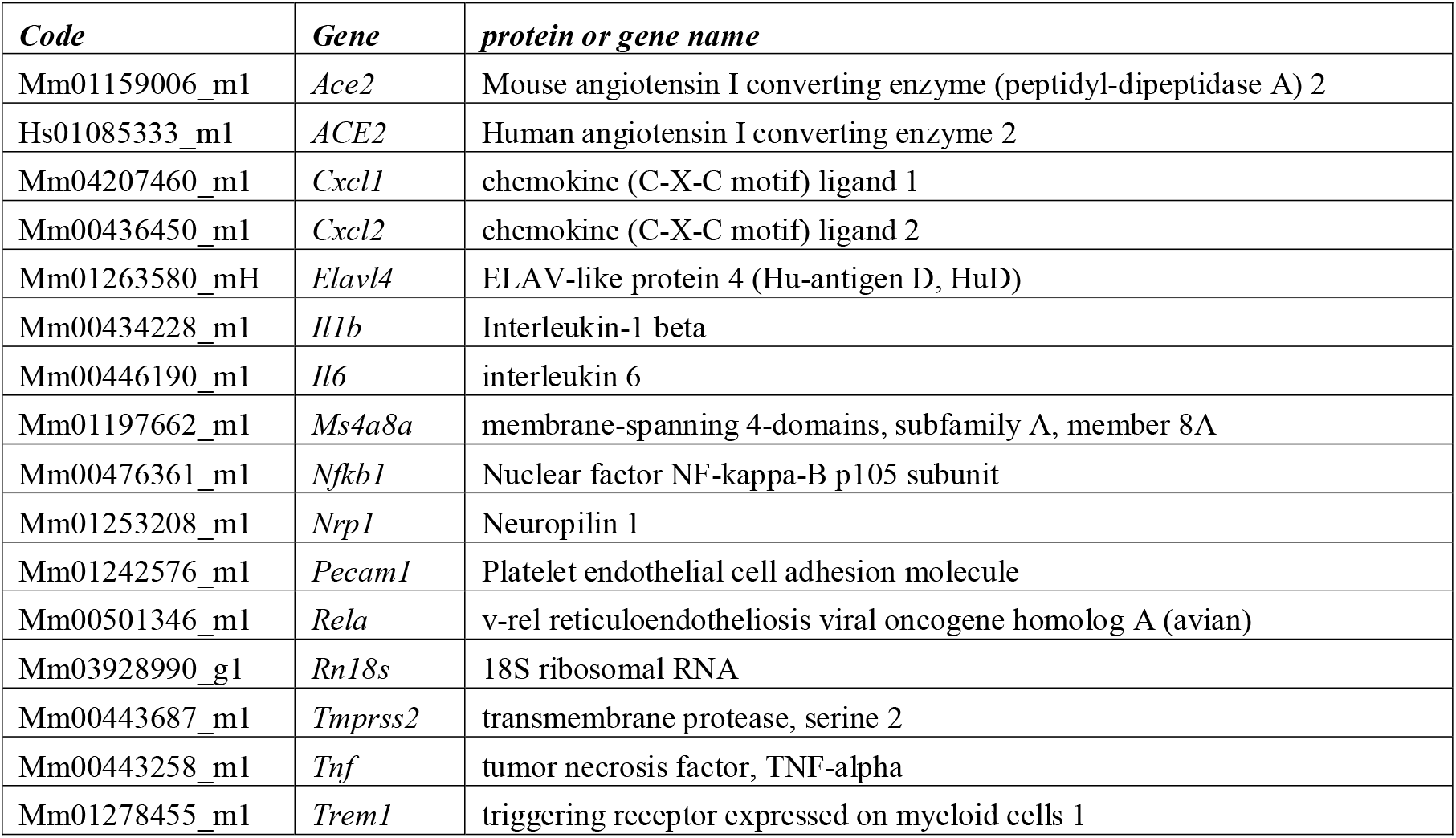
List of TaqMan® probes used in this study.

### Immunofluorescence staining and confocal microscopy

For immunohistochemistry of mouse brain sections, 40 μm-thick floating sections were rinsed 4 times in 0.1 M PBS pH 7.4 and blocked for 30min at room temperature in blocking solution (PBS containing 0.1% BSA, 10% normal donkey serum and 0.4% Triton X-100). Sections were incubated overnight at 4°C with a mix of primary antibodies diluted in blocking solution (see Antibody Table 2). The sections were washed three times in PBS and incubated at room temperature for 2 hours with Alexa Fluor-conjugated secondary antibodies (1:500 dilutions in blocking solution). After three washes with PBS, nuclei were counterstained by incubating the sections for 3 minutes in DAPI (dilution 1:5,000 in PBS) before mounting and coverslip with homemade Mowiol.

**Table 2.** List of antibodies used in this study.

| *Antibody* | *Host* | *Manufacturer* | *Reference* | *Dilution* |
| --- | --- | --- | --- | --- |
| Anti-SARS Cov1/2<br>nucleocapsid protein | mMouse | Sino Biological | 40143-MM05 | 1:200 |
| Anti-mouse | Donkey | Invitrogen | A10037 | 1:500 |
| Anti-rabbit | Donkey | Invitrogen | A21206 | 1:500 |
| Anti-chicken | Donkey | Jackson Immuno<br>Research | 703-605-155 |  |

### Brain vascular pathology analysis

For string vessel measurements, pepsin antigen retrieval was performed for 5 min at 37 °C (0.1 mg ml^−1^ pepsin in PBS, 0.2 N HCl) on 50 μm thick brain sections. For vascular density, an antigen retrieval procedure (20 min at 95°C in 10 mM sodium citrate solution) was performed before the staining. Sections were blocked with 3% BSA in PBS containing 0.1% Triton X-100 for 6h at room temperature, and incubation with primary antibodies (collagen IV: Bio-Rad, #134001, 1:200; caveolin-1: Cell Signaling Technology, #3267, 1:400) was performed at 4 °C overnight, while incubation with secondary antibodies was performed in blocking solution at room temperature for 2h. Images were taken using a confocal laser scanning microscope (Leica, SP5). For all analyses, we imaged four fields from at least two sections per mouse and analysis was performed blinded using ImageJ.

### RNAscope

In situ hybridization was carried out using the protocol from Advanced Cell Diagnostics (ACDbio), following the manufacturer’s guidelines (Cat. No. 323100, RNAscope® Multiplex Fluorescent Reagent Kit v2). To minimize the degradation of the RNA, all the solutions were prepared in Diethyl pyrocarbonate (DEPC, Sigma) treated water (DEPC-H2O). 18-μm thick brain sections, mounted onto SuperFrost Plus Slides (Thermo Fisher), and fixed with PFA 4%, was dehydrated in ethanol 50%, 70% and 100% two times. For hybridization protocol, we followed the manufacturer’s guideline and reagents. Using the probes for mouse Mtnr1a-C3 (cat. No. 403601-C3, lot :21132B), Mtnr1b (cat. No. 502451, lot :21132B) designed by ACDBio. 3-plex negative and 3-plex positive probes were used as negative and positive controls. The positive probe hybridized with mouse Ubc, Ppib and Polr2a mRNA, and the negative probe hybridized with bacterial DapB mRNA (both provided by ACDBio). After RNAscope® assays, a classical immunofluorescence was performed using the Anti-Collagen IV primary antibody (Anti-Collagen IV, Abcam: ab6586, 1:500°) and a secondary Alexa488 Fluor-conjugated antibody (1:500; Molecular Probes, Invitrogen) and DAPI (ACDBio) was used to stain the nuclei. Sections were analyzed using an Axio Imager.Z2 ApoTome microscope (Zeiss, Germany), equipped with a motorized stage and an AxioCam MRm camera (Zeiss, Germany). High-magnification photomicrographs were acquired with a ×63 objective (NA 1.4) using an inverted confocal microscope (LSM 710, Zeiss). Images of target probes and control probes were taken at the same exposure and acquisition parameters.

### Cell culture and transfection

#### HEK293T cells

HEK293T (RRID:CVCL 0063) cells were obtained from Sigma-Aldrich and authenticated by the provider. Cell cultures were maintained in Dulbecco’s Modified Eagle’s Medium (DMEM) Glutamax (Invitrogen) supplemented with 10% fetal bovine serum and 1% streptomycin:penicillin, at 37°C (95% O2, 5% CO2). Cell lines were checked regularly for any mycoplasma contamination.

### hCMEC/D3 cells

For M^pro^-induced cell death assay, hCMEC/D3 cells were cultivated and transfected as described previously ^29^. Briefly, after withdrawing heparin from the medium, we transfected the cells using Lipofectamine 3000 (Thermo Fisher Scientific) and the following plasmids: pCAG-GFP, pCAG-p.Cys145Ala-Mpro-HA or pCAG-Mpro-HA (100 ng per well on 96-well plates). After lipofection, cells were treated with melatonin, AgoMLT, RML (all 100 µM) or DMSO as control for 48h before they were fixed using 4 % PFA for 20 min at room temperature. Staining was performed as described above using primary antibodies against GFP (Abcam, #ab13970, 1:2000) and DAPI (1:2000), and imaged using a fluorescence microscope (DMI6000B, Leica). Analysis was performed blinded using ImageJ.

For pseudovirus luciferase reporter assay, hCMEC/D3 cells were transduced with lentivirus containing hACE2 transcript (BPS Bioscience) at a MOI of 5. Cultures were enriched with positively-transduced cells by antibiotic resistance selection (puromycin resistance gene delivered by the same lentivirus; puromycin at 1 µg/mL, for 7 days), and used as described in the pseudovirus luciferase reporter assay.

### TR-FRET binding assay

Recombinant RBD protein were purchased from SinoBiological (Beijing, China) and reconstituted in water according to provider’s instructions. Fluorescently labelled spike RBD proteins were obtained by custom labelling of the recombinant proteins with d2 fluorophore on lysines with a N-hydroxysuccinimide activated d2 dye in 100mM PO4 buffer (pH8) by Cisbio Bioassays. SNAP-tagged human ACE2 construct was designed as previously reported^37^.

SNAP-tagged ACE2, expressed in HEK293 cells, were fluorescently labelled by incubating cells with a SNAP suicide substrate conjugated to the long-lived fluorophore Terbium cryptate (Tb; Lumi4-Tb, 100 nM; Cisbio Bioassays) in Tag-lite labelling medium (1h, on ice)^64^. After several washes, cells were collected using enzyme-free cell dissociation buffer (Sigma-Aldrich), resuspended in Tag-lite buffer and distributed into a 384-well plate. Cells were pre-incubated (1h) with melatonin at the indicated concentration followed by addition of RBD-d2 (final reaction volume of 14 μl). Melatonin was reconstituted in DMSO at 100 mM stock solution and further dilution were made in either PBS or DMEM culture medium. Final concentration of DMSO was of 0.1% maximum. After 2h incubation (room temperature) TR-FRET signals were detected using a plate reader (Tecan F500; Tecan, Männedorf, Switzerland) with the following settings: excitation at 340 nm (Tb, energy donor), emission at 665 nm (d2, acceptor) and 620 nm (donor); delay of 150 μs; and integration time of 500 μs. Data is expressed as TR-FRET ratio (acceptor/donor). When indicated, TR-FRET ratio was normalized to % of basal or % of control group.

### TR**-**FRET ACE2 conformation assay

ACE2 conformational changes were assessed by performing intramolecular TR-FRET assay, as previously described ^37^. Briefly, Lumi4-Tb-labelled ACE2 cells were incubated with d2-labelled anti-FLAG tag antibody (2 μg/mL, 1h at room temperature; 61FG2DLF, Cisbio Bioassays), followed by addition of non-labelled RBD (5 nM) or melatonin (100 µM) and TR-FRET signal was immediately read during 1h. Data is expressed as delta-TR-FRET ratio, corresponding to the difference in TR-FRET ratio signal between vehicle- and compounds-treated groups.

### In vitro ACE2 binding assay

*In vitro* binding of spike S1 to ACE2 in the absence or presence of melatonin was performed using a commercially available ACE2:SARS-CoV-2 spike S1 inhibitor screening assay kit (BPS Bioscience, San Diego, USA). The assay is based on Streptavidin-HRP detection of spike S1-Biotin protein bound to ACE2-coated wells in a 96-well plate. Compounds were pre-incubated with spike S1-Biotin (10 nM, 1h, RT) and then added to the ACE2-coated wells. After several washes, streptavidin-HRP solution is added (1h, RT) followed by addition of HRP substrate to produce chemiluminescence, which is then measured using a chemiluminescence reader.

### ACE2 enzyme activity assay

The exopeptidase activity of ACE2 was measured using commercially available kit (BPS Bioscience, San Diego, USA). The assay is based on fluorescent detection of a fluorogenic ACE2 substrate, incubated with recombinant ACE2 (1h, RT) in the absence or presence of competitors (melatonin or DX600 (10 µM) competitor as positive control).

### SARS-CoV-2 cell entry assay

SARS-Cov-2 cell entry was assessed using a pseudovirus luciferase reporter assay (BPS Bioscience). The SARS-CoV-2 spike pseudotyped lentivirus are produced with SARS-CoV-2 spike as the envelope glycoproteins. Since the pseudovirions also contain the firefly luciferase gene, the spike-mediated cell entry is monitored via luciferase reporter activity. ACE2-expressing HEK293 cells or hCMEC/D3 cells were pre-incubated with melatonin (1h, 37°C), followed by addition and incubation with the SARS-CoV-2 spike pseudotyped lentivirus (1h, 37°C). Cells were then washed, medium was replaced by OptiMEM medium, and 48h post-transduction the luciferase activity was read by adding luciferase substrate (ONE-Glo EX luciferase assay system, Promega, Wisconsin, USA) using a luminescence plate reader (Envision, Perkin Elmer, Massachusetts, USA).

### Computational chemistry

Molecular system of RBD-ACE2 (6VW1 PDB entry, http://doi.org/10.2210/pdb6VW1/pdb) fully glycosylated with N-glycan pentasaccharides was extracted from the COVID-19 Proteins Library of the CHARMM-GUI Archive (https://doi.org/10.1002/jcc.20945). Docking of melatonin into RBD-ACE2 was performed using Autodock Vina docking engine ^65^ integrated into the UCSF Chimera software ^66^ with an exploration box including the whole RBD molecule-ACE2 complex. Molecular dynamics simulations were computed with GROMACS 2020.5 ^67^. Structural models were hydrated with explicit TIP3P solvent water molecules into a triclinic box of approximately 80 Å x 90 Å x 130 Å. Then, 77 water molecules were substituted by 97 chloride and 123 sodium ions to neutralize the solute with a NaCl solution at 0.16 M. All calculations were performed with the CHARMM36m force field ^68^, and topology of melatonin was carried out from the CGenFF web server ^69^. After harmonic restrains were applied to its α-carbon trace, the system underwent successively 50,000 steps of steepest descent energy minimization, 100 ps of NVT equilibration at 300K with the V-rescale modified Berendsen thermostat, and 100 ps of NPT equilibration with Berendsen barostat. Restraints were left to produce triplicated molecular dynamics trajectories of 500 ns each. Differential analysis of RBD-ACE2 intermolecular interaction between both in complex with or without melatonin was performed with SINAPs tool (Renault et al, J Chem Inf Model, 2021, accepted) dedicated to highlight hydrogen bonds, salt bridges, and aromatic stacking exclusive to one of two trajectories.

### Statistical analysis

Data shown as the means ± SEM. Sample sizes were designed to give statistical power while minimizing animal use. All statistical comparisons were performed using Prism 9 (GraphPad). The specific statistical tests used for each experiment are indicated in the figure legends. One- or Two-way ANOVA with two-stage linear step-up procedure of Benjamini, Krieger and Yekutieli as post-test for multiple comparisons or uncorrected Tukey test was applied and statistical significance was determined as p-value <0.05 (*p<0.05, **p<0.01, ***p<0.001, ****p<0.0001).

## Data availability statements

The authors declare that all data supporting the findings of this study are available within the paper and its supplementary information files.

## ACKNOWLEDGMENTS

We thank all the members of the Jockers lab, Drs Morgane Bomsel, Fernando Real (Institut Cochin) and Francisco Foriestiero (Sao Paulo) for the discussion at the initial phase of the project and Dr. Lara Jehi and her team (Cleveland Clinic) for assistance. We thank Wiebke Brandt and Beate Lembrich (both Lübeck) for expert technical assistance. The SARS-CoV-2 strain *BetaCoV/France/IDF/200107/2020* was kindly provided by Dr JC. Manuguerra, CIBU, Pasteur Institute. The authors thank Dr Pierre Olivier Couraud for the human hCMEC/D3 BBB cell line. The authors thank the Mesocenter of Lille University for computational resources. This work was supported by the Agence Nationale de la Recherche ((ANR-RA-COVID-19 (ANR-20-COV4-0001 to RJ), (ANR-19-CE16-0025-01 to RJ), (ANR-16-CE18-0013 to JD)), Institut National de la Santé et de la Recherche Médicale (INSERM), Centre National de la Recherche Scientifique (CNRS), the Deutsche Forschungsgemeinschaft (WE 6456/1-1 to J.W.). RJ was supported by the Fondation de la Recherche Médicale (Equipe FRM DEQ20130326503) and La Ligue Contre le Cancer N/Ref: RS19/75-127 and E.C. by the Association France Alzheimer (grant No. 2042). JD was supported by the Fondation de France.

## AUTHOR CONTRIBUTIONS

Conceptualization, E.C., M.S., V.P. J.D., and R.J.; In vivo experiments: E.C., C.I, B.K., S.L.P. and J.D. Biochemical investigation: E.C., D.F., C.F.F.C., J.W., S.G.W. and J.D.; Molecular docking and simulations N.R., C.B., E.C.; Data analysis E.C., D.F., C.F.F.C., J.W., J.D., R.J.; Writing – Original Draft, E.C., N.R., J.D. and R.J.; Writing – Review and Editing, E.C., N.R., J.W., M.S., V.P., J.D., R.J.

