## Supplemental figures for "Melatonin drugs inhibit SARS-CoV-2 entry into the brain and virus-induced damage of cerebral small vessels"

### Supplemental Figure S1

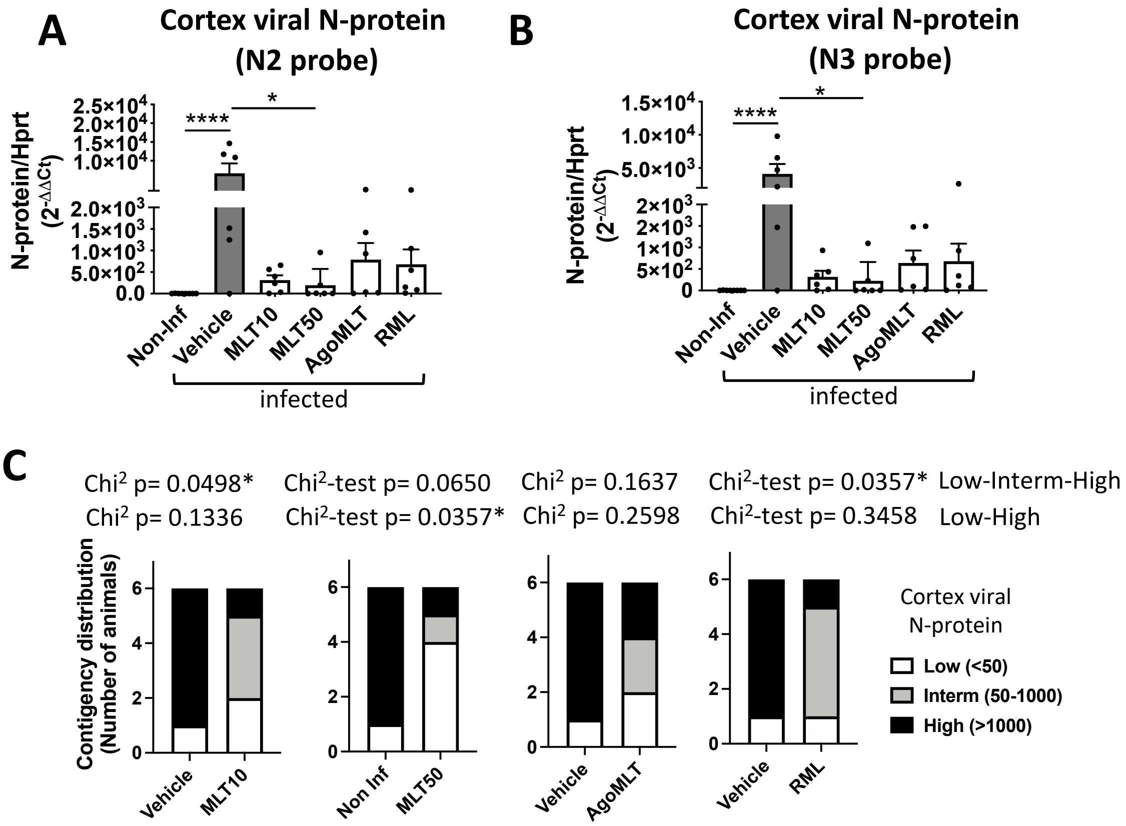


**Supplemental Figure S1 - Melatonergic treatment decreases viral load in the cortex.**
**A-B)** RNA levels of viral N-protein assessed by qRT-PCR with N2 probe (**A**) or N3 probe (**B**) in the cortex of SARS-CoV-2 infected mice at DPI-7. *p<0.05, ****p<0.0001 by Kruskal-Wallis test with two-stage linear step-up procedure of Benjamini, Krieger and Yekutieli as post-test for multiple comparisons. **C.** Contingency distribution of mice grouped in 3 categories per tercentiles according to their viral N-protein RNA level in the cortex (”low”, ”Intermediate”, “High”), for each drug treatment compared to vehicle (Chi$^2$-test, p-value= 0.0498, 0.0650, 0.1637, 0.0357 for MLT10, MLT50, AgoMLT and RML, respectively). The p-values of Chi$^2$-test considering treatment effect versus vehicle for the distribution into “Low” and “High” levels of N-protein is also indicated (p-value=0.1336, 0.0357*, 0.2598, 0.3458 for MLT10, MLT50, AgoMLT and RML, respectively)

#### Supplemental Figure S2

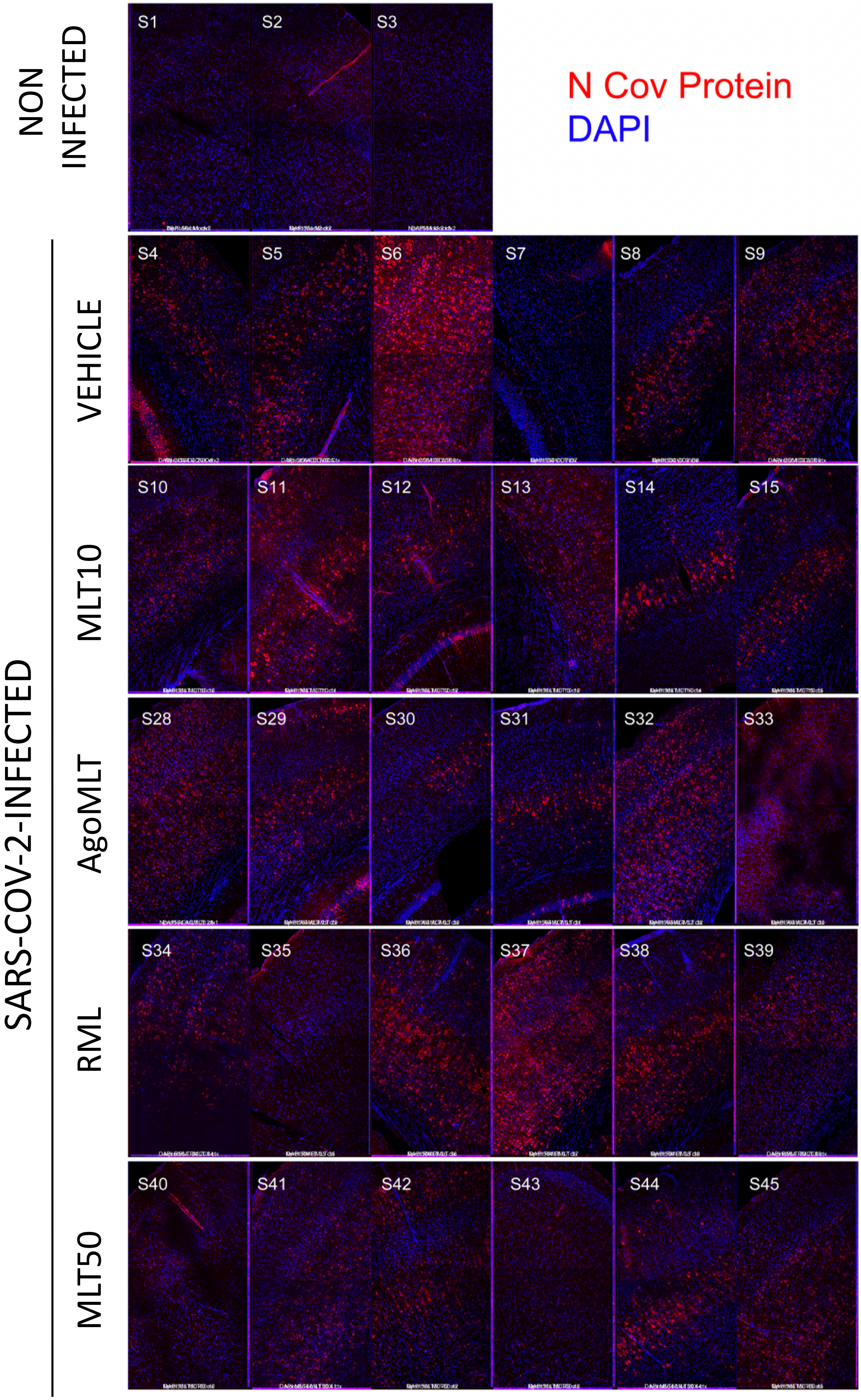


**Supplemental Figure S2 - Viral N-protein in the cortex.** Immunolabeling for N-protein (red) and nucleus (blue) in the cortex of all infected K18-*hACE2* mice 7 days after SARS-CoV2 infection. Individual mice are numbered.

**Supplemental Figure S3**

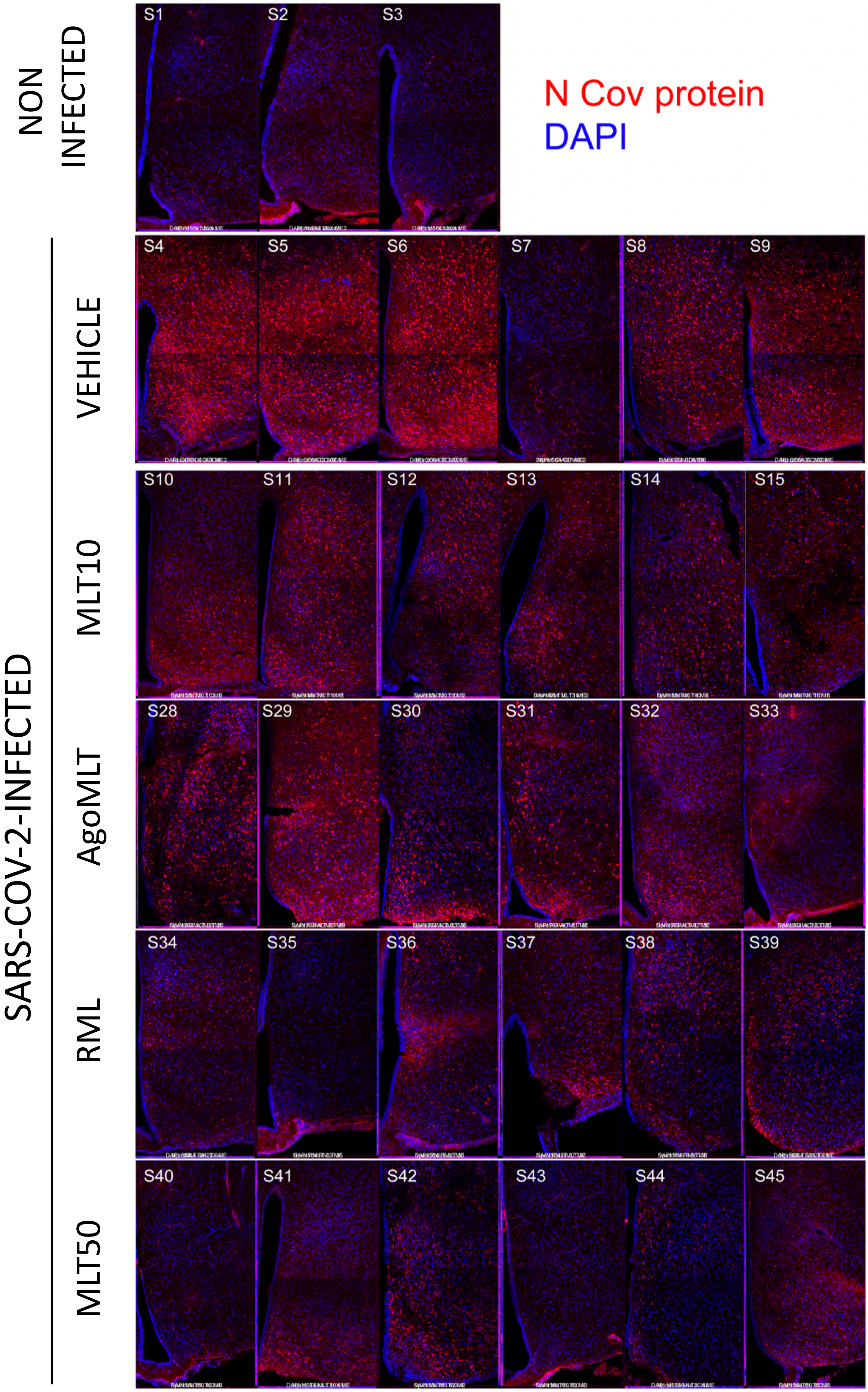


**Supplemental Figure S3 - Viral N-protein in the hypothalamus.** Immunolabeling for N-protein (red) and nucleus (blue) in the hypothalamus of all infected K18-*hACE2* mice 7 days after SARS-CoV2 infection. Individual mice are numbered.

### Supplemental Figure S4

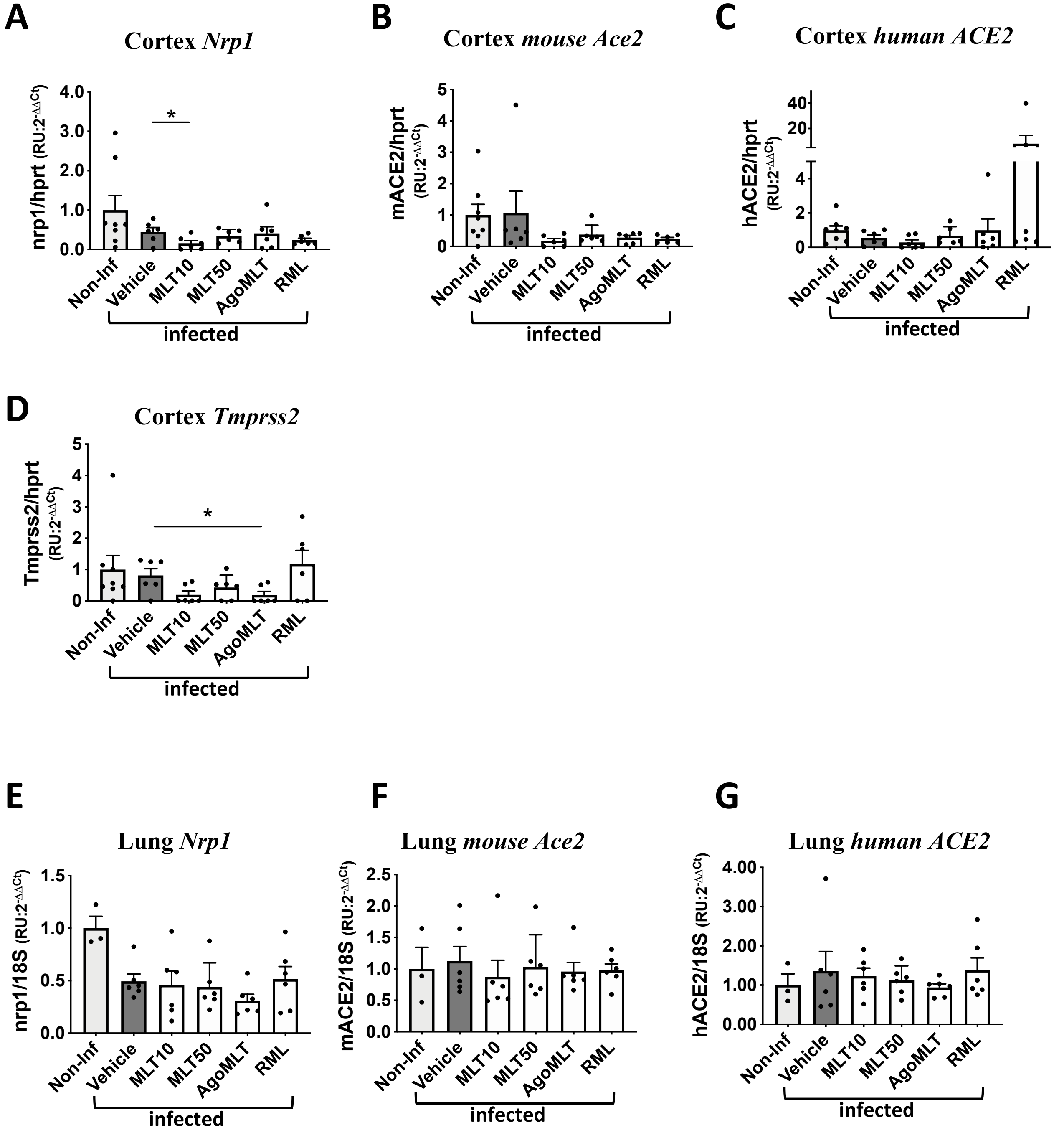


**Supplemental Figure S4 - No effect of compound treatment on the overall expression of *Ace2*, T*MPRSS2* and *NRP1* in the cortex and the lungs. A-E.** mRNA levels of *Nrp1* (**A, E**), mouse *Ace2* (**B,F**), human *ACE2* **(C,G)** and *Tmprss2* **(D)** in cortex (**A-D**) and in the lungs (**E-G**) of SARS-CoV-2 infected mice at sacrifice day 7. Analysis by One-way Anova Kruskal-Wallis test with two-stage linear step-up procedure of Benjamini, Krieger and Yekutieli as post-hoc test for multiple comparisons.

### Supplemental Figure S5

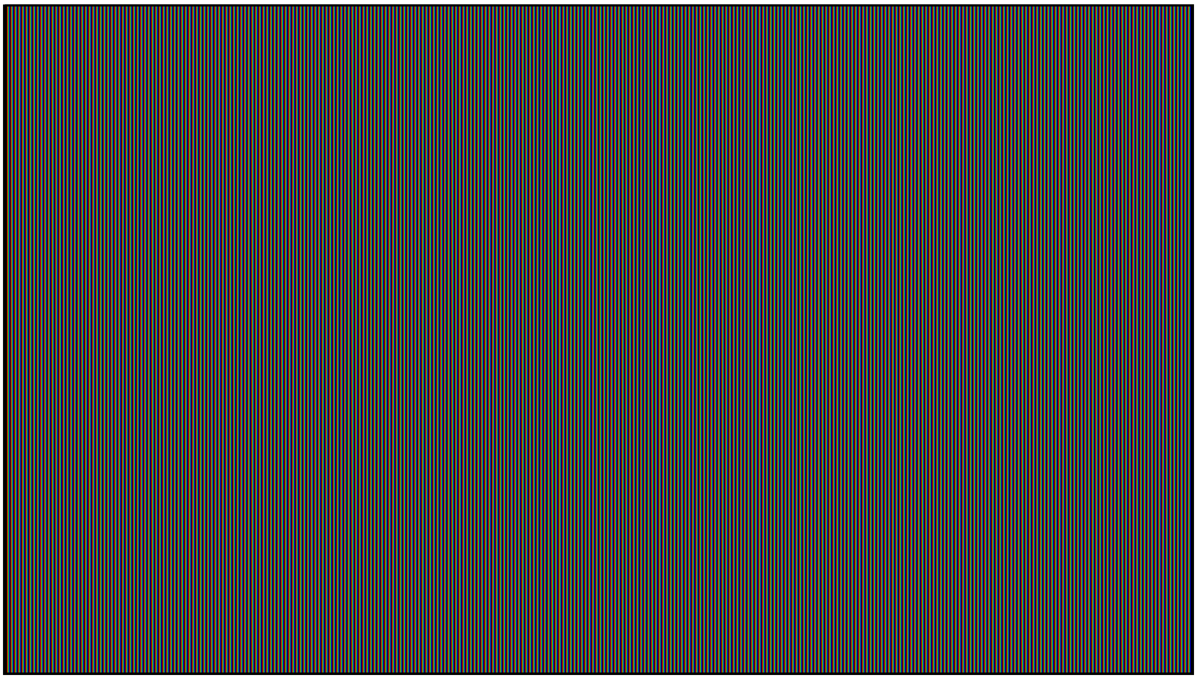

**Supplemental Figure S5 – Animation movie of RBD-ACE2-MLT molecular dynamic. simulation.** Amino acid residues in direct contact with melatonin at t=0 and t=500 nsec of the molecular dynamics simulations are are highlighted.
